# Characterisation of carried and invasive *Neisseria meningitidis* isolates in Shanghai, China from 1950 to 2016: implications for serogroup B vaccine implementation

**DOI:** 10.1101/284133

**Authors:** Mingliang Chen, Charlene M.C. Rodrigues, Odile B Harrison, Chi Zhang, Tian Tan, Jian Chen, Xi Zhang, Min Chen, Martin C.J. Maiden

## Abstract

**Background:** Serogroup B invasive meningococcal disease (IMD) is increasing in China, little is known however, about these meningococci. This study characterises a collection of isolates associated with IMD and carriage in Shanghai and assesses current vaccine strategies.

**Methods:** IMD epidemiological data in Shanghai from 1950–2016 were obtained from the National Notifiable Diseases Registry System, with 460 isolates collected for analysis including, 169 from IMD and 291 from carriage. Serogroup B meningococcal (MenB) vaccine coverage was evaluated using Bexsero® Antigen Sequence Type (BAST).

**Results:** Seven IMD epidemic periods have been observed in Shanghai since 1950, with incidence peaking from February to April. Analyses were divided according to the period of meningococcal polysaccharide vaccine (MPV) introduction: (i) pre-MPV-A, 1965-1980; (ii) post-MPV-A, 1981-2008; and (iii) post-MPV-A+C, 2009-2016. IMD incidence decreased from 55.4/100,000 to 0.71 then to 0.02, and corresponded with shifts from serogroup A ST-5 complex (MenA:cc5) to MenC:cc4821 then MenB:cc4821. MenB IMD became predominant (63.2%) in the post-MPV-A+C period, of which 50% were caused by cc4821, with the highest incidence in infants (0.45/100,000) and a case-fatality rate of 9.5%. IMD was positively correlated with carriage rates. Data indicate that fewer than 25% of MenB isolates in the post-MPV-A+C period may be covered by the vaccines Bexsero®, Trumenba®, or a PorA-based vaccine, NonaMen.

**Conclusions:** A unique IMD epidemiology is found in China, changing periodically from hyperepidemic to low-level endemic disease. MenB IMD now dominates in Shanghai, with isolates harbouring diverse antigenic variants potentially beyond coverage with licenced OMV- and protein-based MenB vaccines.

**Summary:** Meningococcal disease in Shanghai, China is described and current vaccine approaches evaluated. Since 1950, MenA:cc5 shifted to MenC:cc4821 then MenB:cc4821, with MenB dominating since 2009. Distinct antigens potentially beyond coverage with licensed OMV- and protein-based MenB vaccines were found.

## Introduction

*Neisseria meningitidis* is a leading cause of bacterial meningitis and septicaemia globally, with over 1.2 million invasive meningococcal disease (IMD) cases annually [1]. Over 90% of IMD cases are caused by serogroups A, B, C, W, and Y [2], all of which are potentially vaccine-preventable following the licensure of protein-based meningococcal vaccines in 2013 [3].

In China, during the 1950s to 1980s, serogroup A (MenA) isolates were responsible for over 95% of cases [4], with incidence peaking in 1967 (403/100,000) [5]. These were predominantly due to ST-5 clonal complex (cc5) and cc1 [6], and in response, a MenA meningococcal polysaccharide vaccine (MPV) was routinely administered from 1980 onwards [5, 6]. This was followed by a decrease in MenA incidence. From 2003-2005, serogroup C hypervirulent lineage ST-4821 complex (MenC:cc4821) caused outbreaks in Anhui [4], leading to the predominance of MenC IMD and MenC:cc4821 [5, 7]. As a result, in 2008, a serogroup A and C bivalent MPV was introduced into the vaccination program [5], followed by an overall IMD incidence decrease to 0.047/100,000, although this may have been underestimated [6]. From 2011 onwards, the proportion of MenC IMD began to decrease while MenB increased from 7.2% in 2006 to 26.5% in 2014 nationwide [6, 8], with a few regional MenW:cc11 cases [9].

Prevention of MenB IMD is challenging due to the poorly immunogenic polysaccharide capsule and concerns about autoimmunity due to its structural similarity to human tissue. To address this deficit, two protein-based vaccines, Bexsero® (4CMenB) and Trumenba® (bivalent rLP2086), were developed and licensed in Europe and the USA [10, 11]. Bexsero® is composed of factor H binding protein (fHbp), Neisserial heparin-binding antigen (NHBA), *Neisseria* adhesin A (NadA), and PorA, while Trumenba® contains two fHbp-subfamily variants [12]. Both Bexsero® and Trumenba® may elicit protective responses across serogroups [13]. Two methods were established to predict Bexsero® coverage. The Meningococcal Antigen Typing System (MATS) combines phenotypic and functional assays [11]; however, it is time and labour intensive, requires toddler serum, and is only performed by specialist laboratories. Bexsero® Antigen Sequence Typing (BAST) is a rapid, scalable, and portable genotypic approach, which catalogues deduced peptide sequences and matches to vaccine variants (BAST-1) or cross-reactive variants [14].

Limited information is available documenting *N. meningitidis* isolates associated with IMD and carriage in China over the past 60 years. In this study, fluctuations of IMD and meningococcal carriage are described in association with the introduction of MPV vaccines in Shanghai, China since 1950. In addition, and in response to increasing MenB IMD [6], we assessed the potential impact of protein-based vaccines to local prevalent serogroups and clonal complexes.

## Methods

### IMD surveillance

IMD surveillance in Shanghai, implemented in the National Notifiable Diseases Registry System (NNDRS), began in 1950 and was based on monthly paper reports. Since 2004, it has become a web-based, real-time system [5]. All clinical specimens and meningococcal isolates from suspected IMD cases in Shanghai are sent to Shanghai CDC when they are reported in the NNDRS [5]. In China, a child is defined as aged <15 years and an infant <1 year [5].

### *N. meningitidis* carriage surveys

Twenty carriage studies were conducted during 1965-2016. In each study, three districts were chosen, including urban, suburban, and rural districts. Posterior oropharyngeal swabs were collected from preschool children (toddlers aged 3-6 years in childcare centres), students (aged 6-14 years in schools), and adults (staff in department stores, railway stations, army, and residents in communities), and cultured as previously described [15].

### Isolate collection

From 1965-2016, 460 isolates were collected in Shanghai, excluding the period 1986-2004 when isolates were not stored. As a result, 169 IMD and 291 carriage isolates dating from 1965-1985 (n=306) and 2005-2016 (n=154) were available for study. Serogroup was determined by slide agglutination using monoclonal antiserum (BD, USA) and PCR [16]. Isolate serogroup distribution was: A, 123 isolates; B, 221; C, 62; E, 13; W, 5; X, 3; Y, 9; and Z, 3; and 21 nongroupable (NG, negative by PCR and sera agglutination). Sequence type (ST), cc, *porA* and *fetA* variants were determined using PubMLST.org/neisseria [17]. Relationships between STs were analysed using BioNumerics software package (version 7.6.2; Applied Maths, Belgium).

### BAST identification and vaccine coverage estimates

BAST was determined as reported previously [14]. Exact matches and potential cross-reactive matches were combined to evaluate coverage of Bexsero®, Trumenba®, and NonaMen, a 9-valent OMV-based vaccine (Table S1) [18-23].

### Statistical analysis

Statistical analysis was performed using SPSS (version 20.0; IBM, USA). Fisher’s exact test was used to compare proportions of IMD occurring in children with causative serogroup. Statistical significance was assessed at *p* <0.05. The correlation coefficient between carriage rate and IMD incidence was calculated using Microsoft Excel 2010.

## Results

### Epidemiology and characterisation of *N. meningitidis* isolates associated with IMD and carriage in Shanghai

From 1950 to 2016, seven IMD epidemic periods were observed, each lasting 8-10 years (Figure 1A). Average incidence per 100,000 population was: 21 (case-fatality rate, 8.5%) in 1953-1961; 87 (3.3%) in 1962-1972; 2.9 (5.7%) in 1973-1981; 1.6 (6.3%) in 1982-1990; 0.23 (3.7%) in 1991-2000; 0.17 (9.8%) in 2001-2008; and 0.02 (15%) in 2009-2016. Highest incidence occurred in children aged <5 years, decreasing with age, except in those aged 15-19 years from 2005 (Figure 1B). Seasonality of IMD rates was apparent; 50-70% of cases occurred between February and April, with fewer cases (8-23%) between June and October (Figure 1C). A positive correlation was observed between carriage rate and IMD incidence (Figure 2).

**Figure 1.**
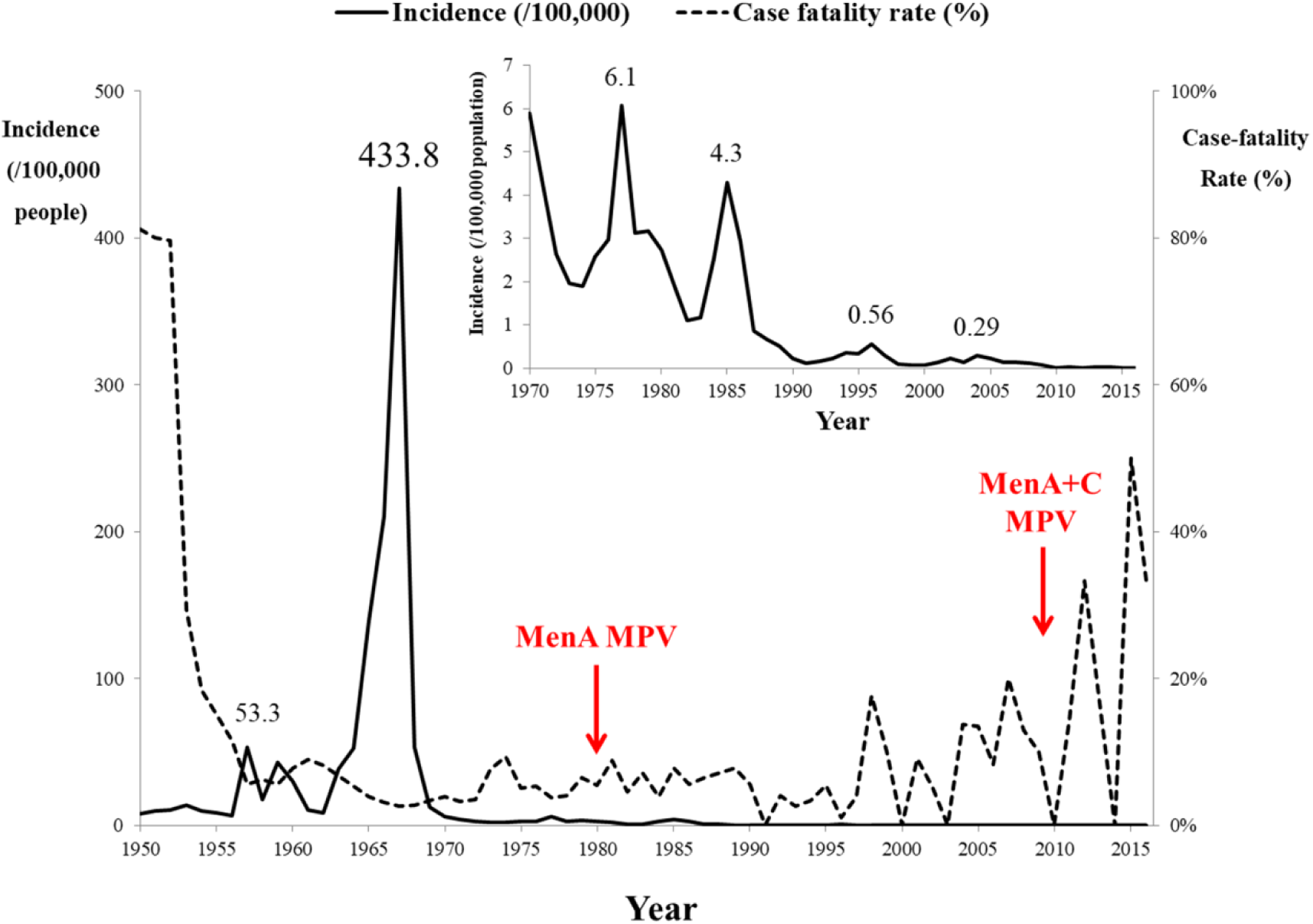
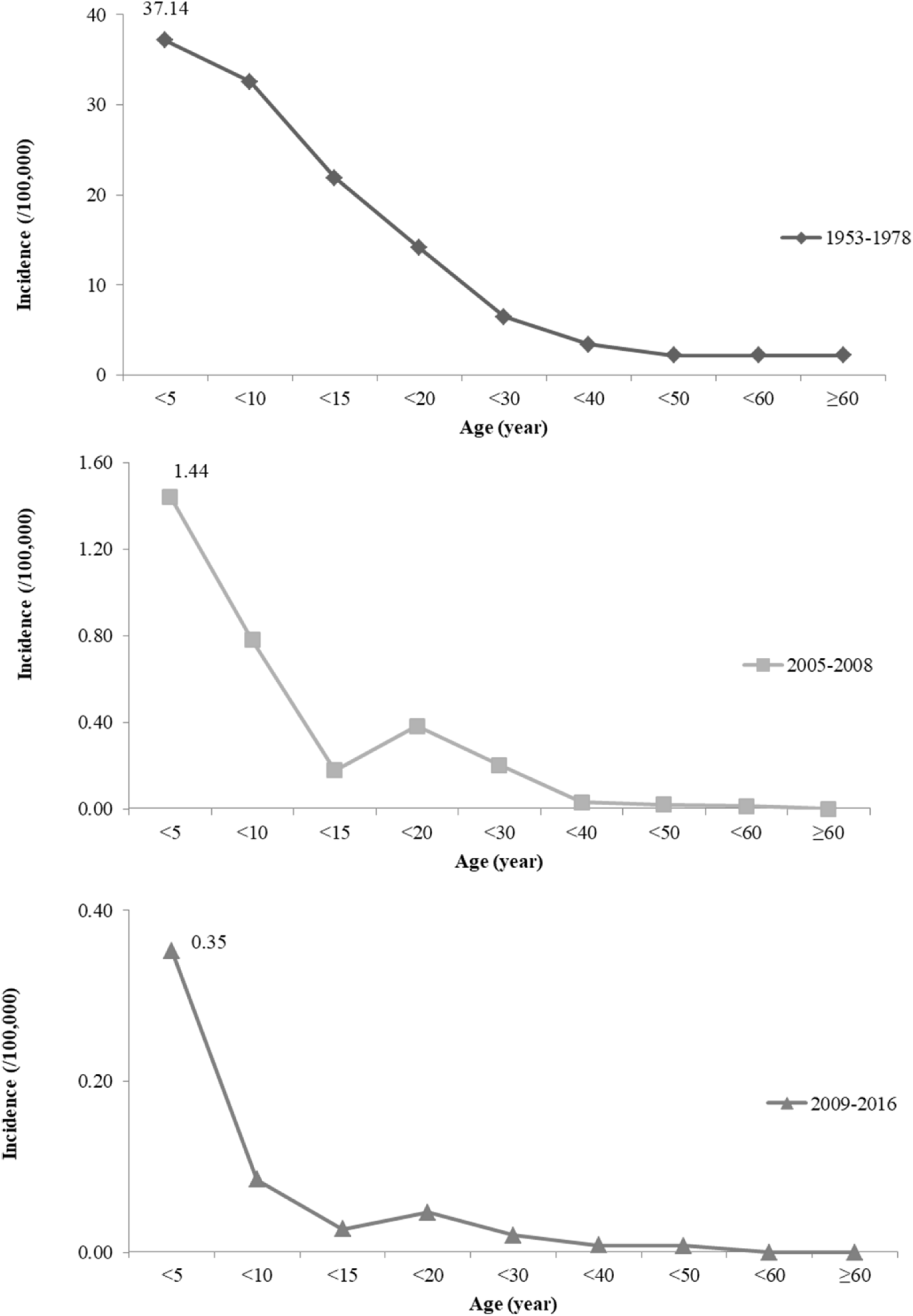
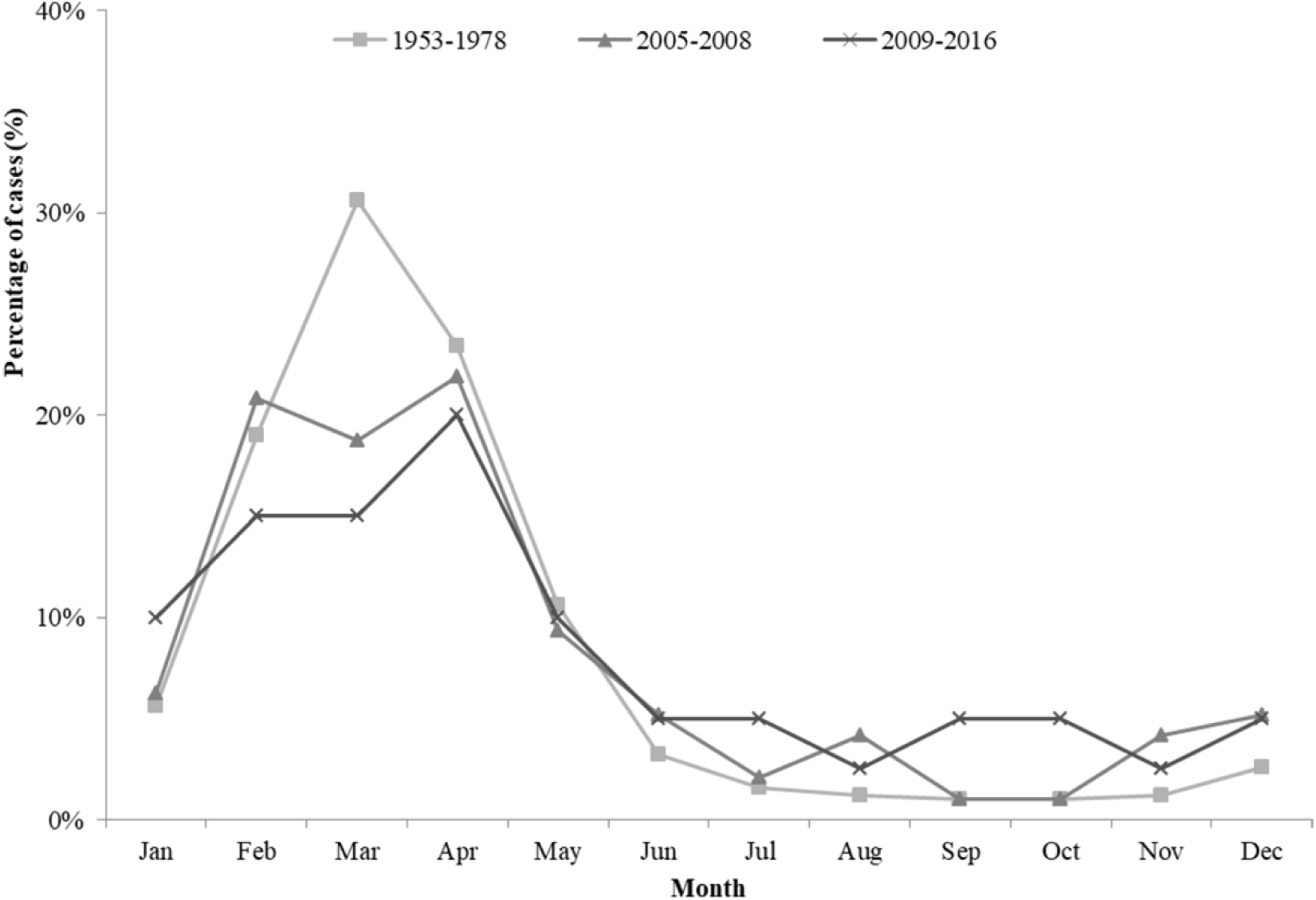
Invasive meningococcal disease incidence in Shanghai, China during 1950-2016, as reported in National Notifiable Diseases Registry System. A) Incidence with case-fatality rates before and after the time of introduction of serogroup A (1980) and serogroups A and C polysaccharide vaccines (2008) in Shanghai, China. Inset figure shows the incidence after 1970. The highest incidences in different epidemic period were labelled. B) Analysis of incidence by age group. C) Seasonality of invasive meningococcal disease in Shanghai, China. MenA, serogroup A meningococcus; MPV, meningococcal polysaccharide vaccine.

**Figure 2.**
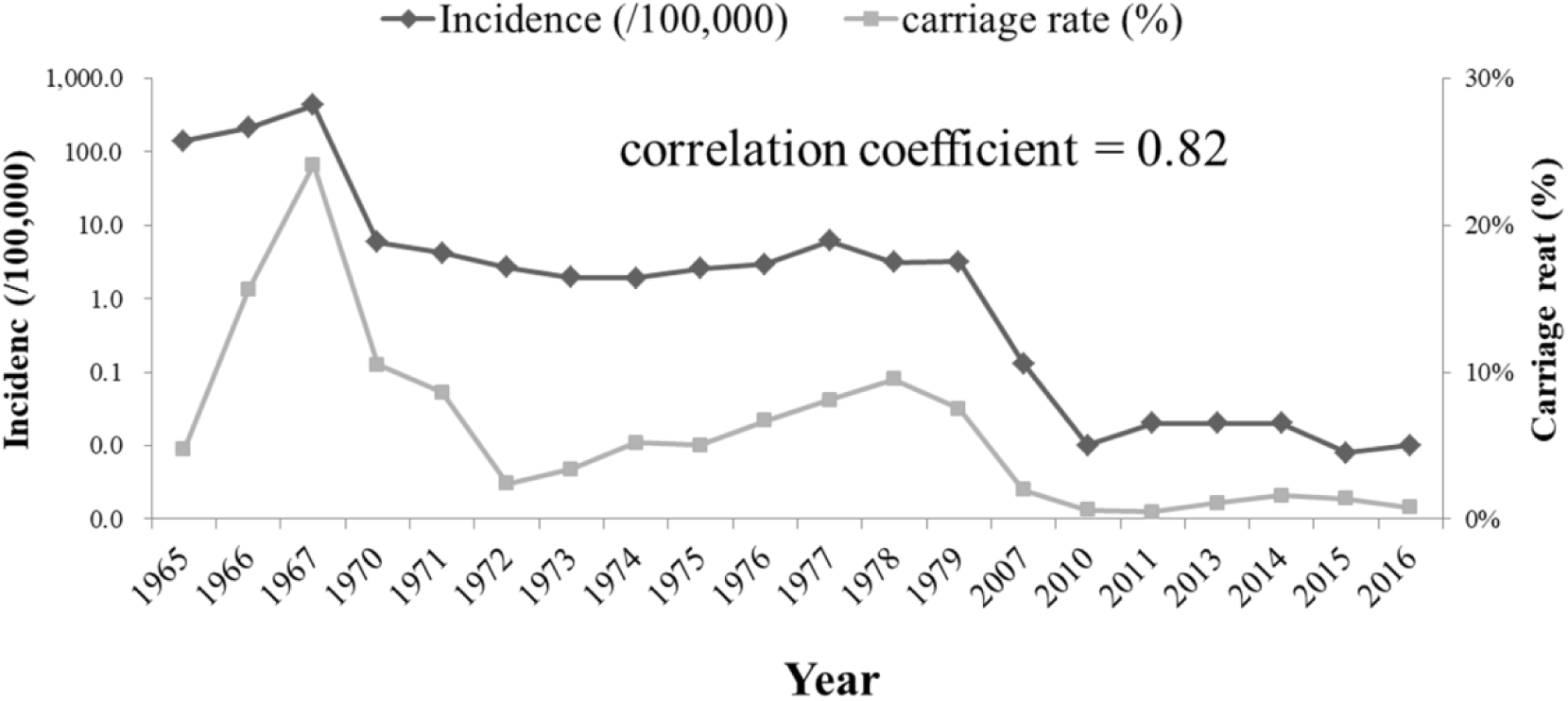
Positive correlation between carriage rate and invasive meningococcal disease incidence in Shanghai, China.

Based on the time of introduction of MPVs in China (1980 and 2008), three periods were defined: (i) pre-MPV-A, 1965-1980; (ii) post-MPV-A, 1981-2008; and (iii) post-MPV-A+C, 2009-2016 (Table 1 and Figure 3).

**Table 1.** Epidemiological information and molecular characterisation of meningococcal isolates before and after introduction of vaccines in Shanghai, China

| Period | Disease isolates |  |  | Carriage isolates |  |  |
| --- | --- | --- | --- | --- | --- | --- |
|  | i) pre-MPV-A §: | ii) post-MPV-A: | iii) post-MPV-A+C: | i) pre-MPV-A: | ii) post-MPV-A: | iii) post-MPV-A+C: |
|  | 1965-1980 | 1981-2008 | 2009-2016 | 1965-1980 | 1981-2008 | 2009-2016 |
|  | (n=117) | (n=61)* | (n=19)† | (n=178) | (n=24) | (n=89) |
| Incidence,<br>/100,000 | 55.4 (range,<br>1.9-433.8) | 0.71 (0.06-4.3) | 0.02 (0.008-0.03) | 9.3% (carriage rate,<br>2,832/30,766) | 2.0%<br>(11/553) | 1.2%<br>(83/6,284) |
| Case fatality<br>rate, % | 3.0<br>(2,918/97,280) | 6.5<br>(168/2,580) | 15<br>(6/40) | NA ¶ | NA | NA |
| Serogroup | A (71.8%, 84)‡, | A (29.5%, 18/61), | A (0%), | A (10.7%, 19), | A (16.7%, 4), | A (0%) |
|  | B (17.1%, 20), | B (24.6%, 15/61), | B (63.2%, 12/19), | B (52.2%, 93), | B (66.7%, 16), | B (84.3%, 75), |
|  | C (7.7%, 9) | C (45.9%, 28/61) | C (36.8%, 7/19), | C (16.9%, 30), | C (4.2%, 1) | C (3.4%, 3), |
| Clonal complex♀ | cc5 (41.0%, 48), | cc4821 (37.5%, 18/48), |  | cc32 (9.6%, 17), |  |  |
|  | cc1 (30.8%, 36) | cc5 (20.8%, 10/48) | cc4821 (75%, 12/16) | cc4821 (8.4%, 15), | cc4821 (29.2%, 7) | cc4821 (25.8%, 23) |
|  |  |  |  | cc5 (7.3%, 13) |  |  |
| PorA VR |  | P1.7-2,14 |  |  |  | P1.21-2,28 (13.5%, |
|  | P1.20,9 (41.0%, | (29.2%, 14/48), | P1.7-2,14 | P1.7-4,13-20 (11.2%, | P1.7-2,14 | 12), P1.22,23-3 |
|  | 48), P1.7-1,10 |  | (43.8%, 7/16) | 20), P1.7,16 (9.0%, | (12.5%, 3), P1.20,9 | (6.7%, 6), |
|  | (29.1%, 34) | P1.20,9 |  | 16), P1.20,9 (7.9%, 14) | (12.5%, , 3) | P1.18-25,9-18 (5.6%, |
|  |  | (20.8%, 10/48) |  |  |  | 5), P1.22,23 (5.6%, 5) |
| FetA VR |  |  |  | F5-8 (11.8%, 21), |  |  |
|  | F3-1 (32.5%, 38), | F3-3 (34.2%, 13/38), |  | F1-7 (10.1%, 18), | F1-5 (12.5%, 3) , | F1-20 (13.5%, 12), |
|  | F5-5 (31.6%, 37), | F3-1 (23.7%, 9/38) | F3-3 (42.9%, 6/14) | F1-15 (9.0%, 16), | F3-1 (12.5%, 3), | F1-91 (10.1%, 9) |
|  |  |  |  | F3-1 (6.7%, 12) | F3-3 (12.5%, 3) |  |
| BAST | 13 (38.5%, 45), | 22 (21.1%, 8/38), | 802 (21.4%, 3/14) | 2300 (9.6%, 17/162) | 22 (12.5%, 3/24) | 2262 (5.6%, 5), |
|  | 794 (29.1%, 34) | 802 (15.8%, 6/38) |  | 13 (6.2%, 11/178) |  | 2433 (4.5%, 4) |

§ MPV-A, serogroup A meningococcal polysaccharide vaccine.
\* 13 isolates or positive DNA not available for multi-locus sequence typing and PorA VR, and another 10 isolates not available for typing of FetA and BAST.
† 3 isolates or positive DNA not available for multi-locus sequence typing and PorA VR, and another 2 isolates not available for typing of FetA and BAST.
¶ NA, not applicable.
‡ The denominator is indicated when it is different from the total number of isolates in this period.
♀ cc1, ST-1 complex; cc5, ST-5 complex; cc32, ST-32 complex; cc4821, ST-4821 complex.

**Table 2.** Comparison of molecular characterisation of ST-4821 complex by serogroup*

| clonal<br>complex | Sequence type | fHbp VR | NHBA VR | PorA VR | PorB VR | FetA VR |
| --- | --- | --- | --- | --- | --- | --- |
| MenB:cc4821<br>(n=40) | ST-5664 (9),<br>ST-5798 (6),<br>ST-3200 (6) | 16 (23) | 669 (13),<br>910 (8) | P1.20,23-x (26) ¶ | 3-229 (12), 3-81 (8),<br>3-460 (5), 3-48 (4) | F1-91 (12), F3-9<br>(6), F5-2 (5) |
| MenC:cc4821<br>(n=32) | ST-4821 (23) | 80 (9), 22 (6),<br>404 (5), 419 (5) | 503 (23) | P1.7-2,14 (15),<br>P1.20,23-x (8) | 3-48 (22) | F3-3 (23) |

\* all cc4821 isolates without *nadA* gene.
¶P1.20,23-x, such as P1.20,23-1 and P1.20,23-3

**Figure 3.**
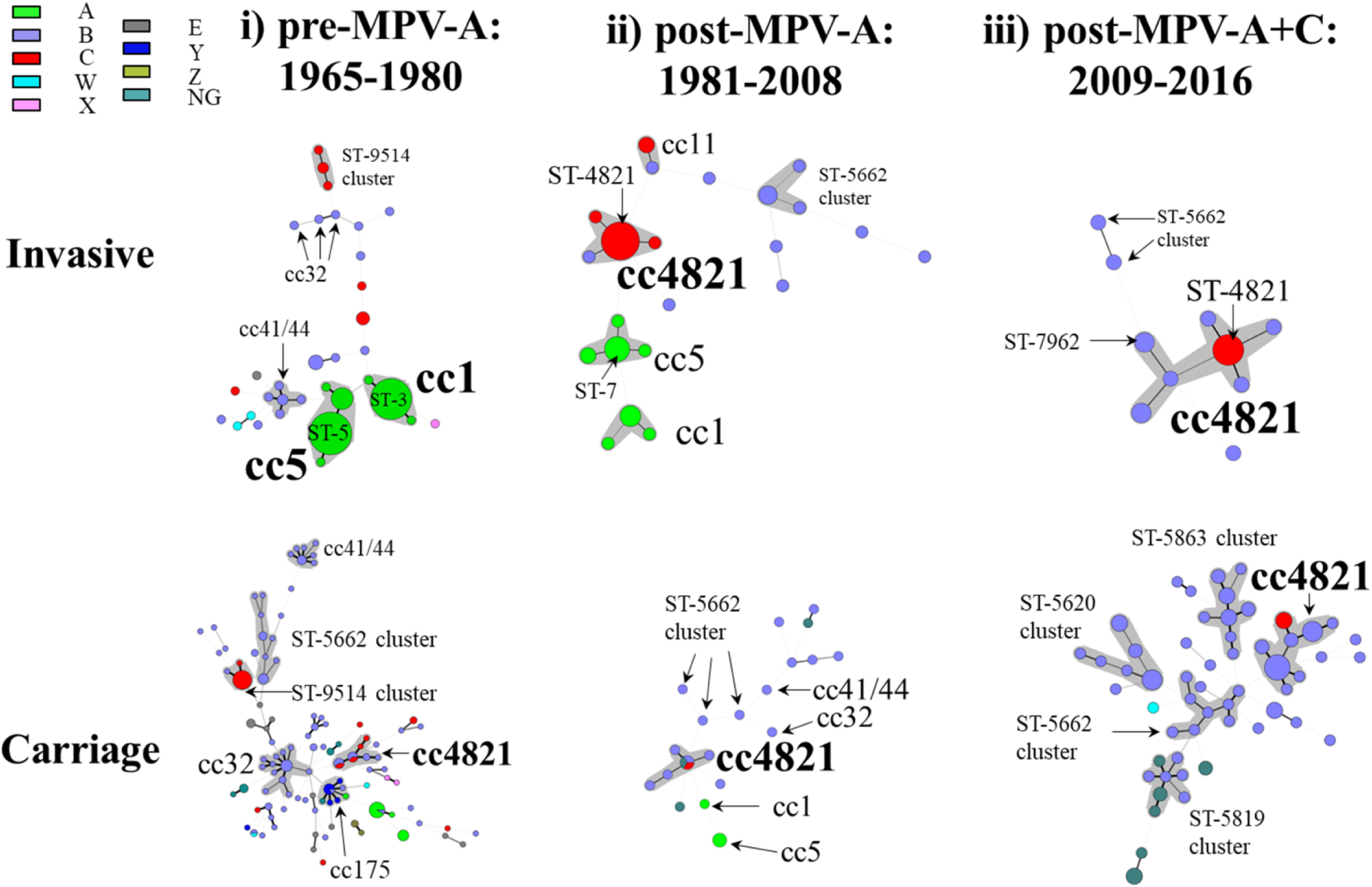
Minimum-spanning tree analysis of multiple-locus sequence types of invasive and carriage *N. meningitidis* before and after introduction of meningococcal vaccines in China. Isolates were obtained during the pre-MPV-A (1965-1980), post-MPV-A (1981-2008), and post-MPV-A+C (2009-2016) periods. Sequence types (STs) are displayed as circles. The size of each circle indicates the number of isolates with this particular type. Serogroup is distinguished by different colours. The shaded halo surrounding the STs encompasses related sequence types that belong to the same clonal complex. Heavy solid lines represent single-locus variants, and light solid lines represent double-locus variants. Sequences types sharing no less than 4 loci, but not assigned to any clonal complexes in the PubMLST database were assigned to ST-clusters. NG, nongroupable.

**Figure 4.**
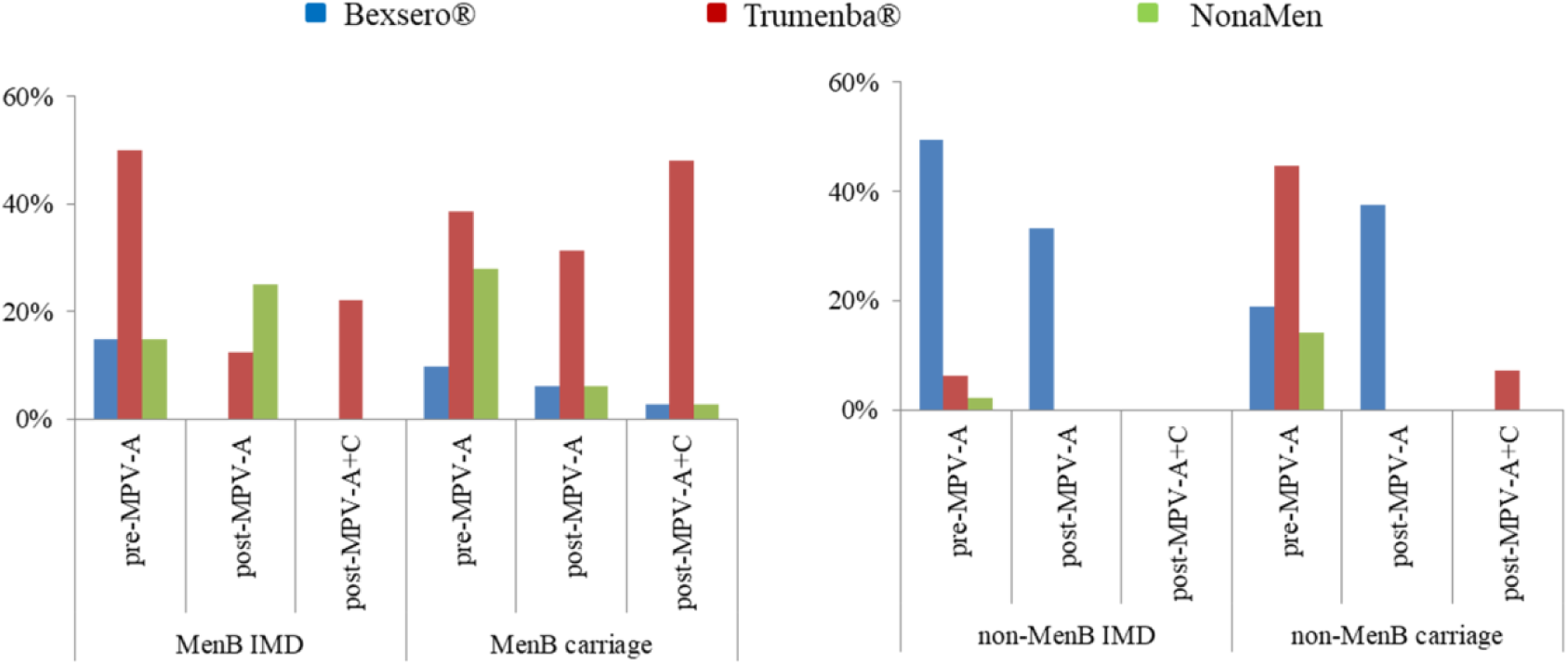
Prevalence of peptide variants, and potentially immunologically cross-reactive variants, for three serogroup B-substitute vaccines, Bexsero®, Trumenba®, and NonaMen, among 460 invasive and carriage meningococci from Shanghai, China in the pre-MPV-A, post-MPV-A, and post-MPV-A+C periods. Bexsero® and Trumenba® are two protein-based serogroup B substitute meningococcal vaccines, which have been licensed in Europe and the USA, while NonaMen is a 9-valent investigational outer membrane vesicle vaccine, which has undergone pre-clinical testing. Three periods were defined, pre-MPV-A (1965-1980), post-MPV-A (1981-2008), and post-MPV-A+C (2009-2016), according to the time of two meningococcal polysaccharide vaccines introduced in China (1980 serogroup A, 2008 A and C).

(i) In the pre-MPV-A period, the average incidence was 55.4/100,000. MenA isolates were predominant (71.8%, 84/117; Table 1), belonging to cc5 (57.1%, 48/84) and cc1 (42.9%, 36/84). Among MenA:cc5 isolates, ST-5 prevailed (77.1%, 37/48), with no ST-7, and all contained PorA VR P1.20,9, while of the MenA:cc1 isolates, 34/36 (94.4%) were ST-3, with 32/34 (94.1%) P1.7-1,10. MenB isolates were assigned to cc41/44 (30%, 6/20), cc32 (15%, 3/20), cc8 (5%, 1/20), cc35 (5%, 1/20), and cc198 (5%, 1/20), with 8 singletons. MenC isolates were assigned to ST-9514 cluster (44.4%, 4/9), cc4821 (33.3%, 3/9), and cc231 (11.1%, 1/9). MenC:cc4821 isolates were all ST-3436 with P1.20-3,23-x, such as P1.20-3,23-1 and P1.20-3,23-3.

The carriage rate ranged from 2.4% in 1972 to 24.1% in 1967 (Table S2), with overall carriage rates of 4.4% (368/8,319) in children and 9.9% (888/8,956) in adults (≥15 years). In 1966-1967, high IMD incidence (>200/100,000) coincided with high carriage rates (>15%), of which a high proportion (>70%) was MenA. This decreased from 50% in 1970 to 1.1% in 1979. Among the 178 carriage isolates analysed, MenB (52.2%) was predominant (Table 1), with cc32 (18.3%, 17/93) the most prevalent.

(ii) In the post-MPV-A period, the average incidence was 0.71/100,000. Based on 61 IMD cases with available serogroup data, MenC (45.9%, 28/61) was the most frequent, in which isolates belonging to cc4821 (89.5%, 17/19) dominated with the majority of these ST-4821 (88.2%, 15/17) and P1.7-2,14. MenA:cc5 (62.5%, 10/16) dominated in MenA isolates, with 8 were collected during 2005-2008 with 6/8 (75%) ST-7 and P1.20,9, and 2 from 1985 ST-5, P1.20,9. MenB isolates were assigned to cc4821 (14.3%, 1/7; ST-5798 with P1.10,13-1), cc41/44 (14.3%, 1/7), and cc32 (14.3%, 1/7), with 4 singletons.

The carriage rate from the 2007 survey was 2.0% (11/553), with 2.4% (9/369) of this in children and 1.1% (2/184) in adults (15-46 years). MenB (66.7%, 16/24) was predominant in carriage (Table 1), 31.3% of which belonged to cc4821, with 5 different STs each possessing a different PorA VR type.

(iii) In the post-MPV-A+C period, the average incidence was 0.02/100,000. MenB (63.2%) isolates predominated, 50% of which were cc4821 and assigned to 5 STs each with a different PorA VR type (Figure 3). All 7 MenC isolates were assigned to cc4821. Except one DNA sample with incomplete ST, other 6 MenC:cc4821 isolates were ST-4821, with 5 containing P1.7-2,14.

The carriage rate ranged from 0.5% in 2011 to 1.6% in 2014, with 1.5% (25/1,660) in children and 1.6% (73/4,624) in adults (15-78 years). MenB (84.3%, 75/89) was the most frequent serogroup in carriage (Table 1), with 20/75 (26.7%) cc4821.

### Features and seasonality of MenB IMD

From 1965 to 2016, 72.3% (34/47) of all MenB IMD occurred in children, while only 40% (60/150) of non-B IMD cases occurred in this age group (p=0.01). Since 2005, all MenB IMD cases were in children (19 days to 12 years), among which 65% (13/20) were infants. During 2005-2008, MenB IMD incidence was 0.01/100,000, highest among infants (1.1/100,000) compared to 0.009/100,000 in children aged 1-15 years, with no reported deaths. During the post-MPV-A+C period, MenB IMD incidence was 0.007/100,000, the highest of which in infants (0.45/100,000) compared to 0.03/100,000 in children aged 1-15 years, with a case-fatality rate of 9.5% (2/21). During the post-MPV-A+C period, MenB cases were observed from February to September, and in December while all MenB cases from 2005-2008 occurred from January to June.

### BAST identification, prevalence of vaccine antigens and potentially cross-reactive variants

A total of 243 BASTs were identified with high diversity in each of the vaccine antigens: fHbp, 64 variants; NHBA, 95; NadA, 9, the *nadA* gene was absent or had gene-silencing frameshift mutations in 82.0% (367/460) of isolates; PorA VR1, 38; and PorA VR2, 64.

A total of 56 BASTs were identified in the 169 IMD isolates (0.33 BASTs/isolates). The four most prevalent BASTs were BAST-13 (cc5), BAST-794 (cc1), BAST-802 (cc4821) and BAST-22 (cc5), represented by 60.4% (102/169) isolates. In the 291 carriage isolates, 201 BASTs were identified (0.69 BASTs/isolates). The four most prevalent BASTs, including BAST-2300 (ST-9514 cluster), BAST-13 (cc5), BAST-794 (cc1), and BAST-2262 (ST-5620 cluster), were represented by 15.0% (40/267) isolates. BASTs fluctuated with ccs found in the pre- and post-MPV periods (Table 1).

Combined exact matches and putative cross-reactive antigens, revealed that 6.8% (15/221) of MenB isolates were potentially covered by Bexsero®, and among IMD MenB isolates, these constituted: 15% (3/20) in pre-MPV-A; 0% in post-MPV-A; and 0% in post-MPV-A+C periods. For Trumenba®, no exact antigen match was found and putative cross-reactive variants were 90/221 (40.7%) among MenB isolates. In IMD MenB isolates this constituted: 50% (10/20), 12.5% (1/8), and 22.2% (2/9) in each respective period. For NonaMen, the covered antigen in MenB isolates was 34/221 (15.4%), and in IMD MenB isolates, the prevalence was 15% (3/20), 25% (2/8), and 0% respectively (Figure 5).

**Figure 5.**
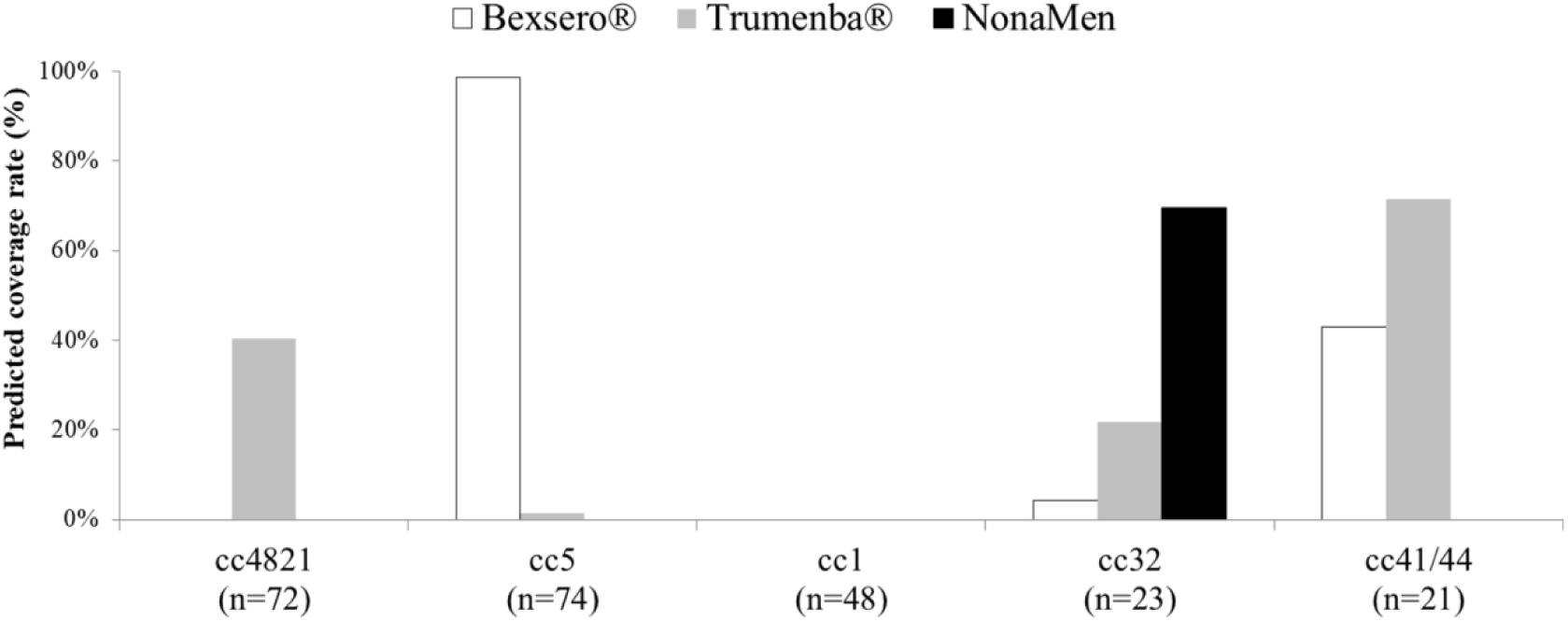
Potential coverage of three serogroup B vaccines, Bexsero®, Trumenba®, and NonaMen to the 5 most prevalent clonal complexes (cc) in Shanghai. For Bexsero®, the prevalence of potentially covered variants was low: cc1, 0%; cc4821, 0%; cc32, 4.3% (1/23); and cc41/44, 42.9% (9/21), except cc5 (98.6%, 73/74). For Trumenba®, the potentially covered antigens were: cc32, 21.7% (5/23); cc4821, 40.3% (29/72); cc41/44, 71.4% (15/21); cc1 and cc5 < 2%. For NonaMen, no antigens were observed in isolates from cc1, cc5, cc41/44 or cc4821, while 69.6% (16/23) of cc32 isolates contained homologous PorA sequences.

## Discussion

This study provides a comprehensive analysis of IMD in Shanghai, China, comparing invasive meningococci with those obtained from carriage. From 1965 to 1980, IMD was dominated by MenA isolates belonging to cc5 (ST-5) and cc1 (ST-3), resulting in several epidemics (Figure 1A and 3). This was consistent with that seen elsewhere with MenA:cc5 meningococci responsible for the first and second pandemic waves between the 1960s and 1990s [24, 25]. Introduction of serogroup A MPV vaccine in China in 1980 was followed by a decrease in IMD; however, this in turn may have contributed to expansion of MenC IMD caused by MenC:cc4821 [4]. The pattern of clonal expansion following vaccine implementation was further observed with the subsequent implementation of serogroup A and C MPV in 2008 which was followed by an increase in MenB IMD, largely due to MenB:cc4821 isolates (Figure 1A and 3) [26]. These data indicate that vaccine intervention may have facilitated the emergence of new strains not targeted by the vaccines, consistent with the secular fluctuation of hyperinvasive lineages. Indeed, similar changes subsequent to vaccine implementation with MPV A+C were observed in Egypt and Morocco during 1992-1995 [27]. This epidemiology appears to be unique to China [6], with a dramatic shift from hyperepidemic disease (55.4/100,000 in 1965-1980), similar to that of low-income regions, to low incidence endemic disease (0.02/100,000 in 2009-2016), more similar to epidemiology of industrialised regions.

Since the 1950s, the seasonal peak of IMD cases in Shanghai has been from February to April (Figure 1C), identical to that seen nationwide [5]. China is, however, a large country, with notable differences seen for example in the peak influenza season between northern (January) and southern China (from June to July), with the latter warmer and more humid [28]. This suggests that, besides climate and influenza incidence, social factors including mass gathering events should be considered when deploying preventative strategies. IMD outbreaks, such as the MenA:cc5 global pandemic and MenW:cc11 transmission, are often associated with the movement of large numbers of people [24, 29]. Similarly, the 1967 MenA epidemics across China (403/100,000) occurred following the National Great Networking event during 1966-1967 [5, 30], where millions of students from all over the country gathered [30]. Correspondingly, the seasonal IMD peaks observed (Figure 1C) may be associated with the Spring Festival, the Chinese New Year. Annually, from January to March, over 200 million people embrace the “Spring Festive travel rush” traveling across the country by train [31], to gather with family and friends. Poor sanitary conditions and overcrowded environments on public transport will facilitate transmission of meningococci. Such information should be considered for optimal future vaccination strategies, to prevent transmission resulting from travel and social gatherings. In addition, these data indicate that more research into meningococcal carriage before, during, and after the Chinese New Year is required.

Indeed, few carriage surveys in China have been undertaken; however, two studies in the Shandong and Guangxi provinces identified high carriage rates of meningococci from hyperinvasive ccs in association with IMD outbreaks [9, 32]. This is consistent with results from our study where carriage rates positively correlated with incidence (Figure 2), with MenB predominant in carriage both pre- and post- introduction of MPVs (Table 1), and cc4821 increasing from 8.4% in pre-MPV-A to 29.2% in post-MPV-A subsequently stabilizing at 25.8% in post-MPV-A+C periods (Table 1). In addition, MenB cases were not linked to a distinct seasonal pattern. Since the 1950s, IMD cases in Shanghai predominantly occurred from February to April (Figure 1C), while during the post-MPV-A+C period, MenB IMD cases occurred more consistently throughout the year. Since the IMD diagnostic criteria require the onset of disease during the epidemic season [5], results from this study indicate that diagnostic criteria should be redefined so that future IMD cases can be accurately diagnosed and reported.

To our knowledge, this study was the first to assess coverage with licensed meningococcal vaccines. Since the 1980s, three monovalent OMV-based MenB vaccines have been licensed for IMD epidemics but they demonstrated clinical efficacy only against homologous meningococci [12]. Although a nonavalent OMV-based MenB vaccine has been evaluated [18], we found low prevalence of its homologous variants among Chinese MenB meningococci based on PorA data in this study (<5%) and from 27 provinces of China (<11%) [26]. Two protein-based MenB substitute vaccines were licensed and implemented in vaccination interventions in Europe and the USA [10, 11]. The coverage of MenB isolates by Bexsero® in the UK during 2014/15 was predicted to be 60.8% using BAST [14], and 66% by MATS [33]. For Trumenba®, coverage rates of 78-100% to collections of diverse strains in Europe and the USA was estimated using serum bactericidal assay [20-23]. In this study, the presence of potentially covered variants was low in Shanghai for both Bexsero® (≤15%) and Trumenba® (≤50%) and, based on fHbp data from 30 provinces across China [34], Trumenba® was predicted to potentially cover 32.5% of IMD and 40% of MenB carriage isolates. The low prevalence is attributed to the predominant cc, cc4821 [26], which has a low prevalence of homologous antigens to Bexsero® (0%) and Trumenba® (40.3%) (Figure 5). In Europe and the USA, MenB cases are mainly due to cc32, cc41/44, and cc269 meningococci [35], which exhibited different antigenic profiles in China. Chinese cc32 isolates contained fHbp peptide 101 (56.5%), which was not present in Bexsero®, while cc32 in Europe predominantly expressed fHbp peptide 1 [35], the Bexsero® variant. Chinese cc41/44 isolates mainly harboured fHbp peptide 19 (71.4%) and PorA-VR2 variant 25 (33.3%), while European cc41/44 meningococci mostly include fHbp peptide 1 and PorA-VR2 variant 4 [35], part of the Bexsero® vaccine. Therefore, the likely impact of Bexsero® on Chinese cc32 (4.3%) and cc41/44 (42.9%) was lower than in Europe (93-100%) [33]. Alternative approaches include an OMV-based vaccine specific for MenB cc4821, and the characterisation of ST and antigen data, especially PorA variants, reported here will be invaluable in assessing vaccine coverage and future serogroup B-substitute vaccine development in China.

Although data in this study are limited by incomplete records of MenB IMD cases during 1950-2004 and the small number of isolates collected during 1965-1980 and 1981-2008; data from children’s hospitals in Shanghai and Beijing provide some insight into MenB IMD during 1976-2002 [36-38], where the two features of MenB IMD, the occurrence in young children and lack of seasonal variation, have persisted since the 1970s. Our findings show that MenB has dominated IMD in Shanghai since 2009. At the time of writing, cc4821 isolates were the predominant cause of MenC and MenB IMD across 27 provinces in China [26]. Besides Shanghai, MenB:cc4821 were also found in 18 other provinces, with two IMD-associated MenB:ST-4821 isolates discovered clustering with MenC:ST-4821 outbreak isolates by genomic analysis [39]. These MenC:cc4821 outbreak strains accounted for the increase from <5% before 2003 to 58% during 2003-2008, resulting in an increase in IMD incidence due to MenC from 0.11 in 2000 to >5.5 during 2004-2007 per 100,000 in Hefei, China [7, 40]. Enhanced surveillance of IMD is therefore essential to monitor changes in cc and antigenic variants of MenB IMD, through vaccine selective pressure or secular change.

Our data suggest that vaccine coverage of MenB:cc4821 meningococci by licensed OMV- and protein-based MenB vaccines may be limited.. Therefore a cautious, region-specific approach to implementation of new protein-based meningococcal vaccines should be considered. Further, the temporal analysis suggests that vaccine implementation coinciding with the start of the calendar year, so as to disrupt transmission events during Spring Festival could have a higher impact. In conclusion, our data indicate that IMD surveillance should be enhanced, combined with comprehensive carriage studies to assess the impact of vaccines in inducing herd immunity.

