## Supplementary Materials for "Characterisation of carried and invasive *Neisseria meningitidis* isolates in Shanghai, China from 1950 to 2016: implications for serogroup B vaccine implementation"

**Supplementary Material**

**Table S1. Variants used to assess the strain coverage of serogroup B vaccines**

| Vaccines | Exact matches | cross-reactive matches | References |
| --- | --- | --- | --- |
| Bexsero® | fHbp variant 1, NHBA variant 2, NadA variant 8, or PorA-VR2: 4 | fHbp variants: 1, 4, 13, 14, 15, 37, or 232, or NadA family: 1 or 2/3 | [1] |
| Trumenba® | fHbp variants: 45 or 55 | fHbp variants:1, 4, 13, 14, 15, 16, 19, 21, 23, 24, 25, 30, 47, 76, 87, 180, 187, 252, 276, or 510 | [2-5] |
| NonaMen | PorA variants: P1.7,16, P1.5-1,2-2, P1.19,15-1, P1.5-2,10, P1.12-1,13, P1.7-2,4, P1.22,14, P1.7-1,1 and P1.18-1,3,6 | NA* | [6] |

* NA, not applicable.

**Table S2. Analysis of 20 surveys of meningococcal carriage in Shanghai from 1965 to 2016.**

| Year | Incidence (/100,000) | Number of throat swabs | Isolates | Serogroup (%) | | | | |
| --- | --- | --- | --- | --- | --- | --- | --- | --- |
|  |  |  | (carriage rate, %) | Isolates tested for serogroup | A | B | C | Other |
| 1965 | 139.5 | 5,367 | 253 (4.7) | 0 | ND* | ND | ND | ND |
| 1966 | 210.7 | 249 | 39 (15.6) | 13 | 10 (76.9) | 1 (7.7) | 1 (7.7) | 1 (7.7) |
| 1967 | 433.8 | 5,590 | 1,347 (24.1) | 259 | 188 (72.6) | 44 (17.0) | 21 (8.1) | 6 (2.3) |
| 1970 | 5.9 | 818 | 86 (10.5) | 12 | 6 (50) | 0 | 6 (50) | 0 |
| 1971 | 4.2 | 897 | 77 (8.6) | 9 | 5 (55.6) | 0 | 4 (44.4) | 0 |
| 1972 | 2.7 | 4,639 | 111 (2.4) | 71 | 10 (14.1) | 43 (60.6) | 18 (25.3) | 0 |
| 1973 | 2.0 | 2,025 | 69 (3.4) | 22 | 3 (13.6) | 19 (86.4) | 0 | 0 |
| 1974 | 1.9 | 1,299 | 68 (5.2) | 0 | ND | ND | ND | ND |
| 1975 | 2.6 | 855 | 43 (5.0) | 10 | 0 | 10 (100) | 0 | 0 |
| 1976 | 3.0 | 779 | 52 (6.7) | 35 | 0 | 29 (82.9) | 6 (17.1) | 0 |
| 1977 | 6.1 | 3,561 | 289 (8.1) | 223 | 3 (1.3) | 196 (87.9) | 18 (8.1) | 6 (2.7) |
| 1978 | 3.1 | 2,326 | 221 (9.5) | 184 | 1 (0.5) | 146 (79.3) | 11 (6.0) | 26 (14.1) |
| 1979 | 3.2 | 2,361 | 177 (7.5) | 177 | 2 (1.1) | 123 (69.5) | 12 (6.8) | 40 (22.6) |
| 2007 | 0.13 | 553 | 11 (2.0) | 11 | 0 | 11 (100) | 0 | 0 |
| 2010 | 0.01 | 644 | 4 (0.6) | 4 | 0 | 2 (50) | 0 | 2 (50) |
| 2011 | 0.02 | 210 | 1 (0.5) | 1 | 0 | 1 (100) | 0 | 0 |
| 2013 | 0.02 | 1,500 | 17 (1.1) | 17 | 0 | 16 (94.1) | 0 | 1 (5.9) |
| 2014 | 0.02 | 3,330 | 54 (1.6) | 54 | 0 | 45 (83.3) | 2 (3.7) | 5 (9.3) |
| 2015 | 0.008 | 360 | 5 (1.4) | 5 | 0 | 3 (60) | 0 | 2 (40) |
| 2016 | 0.01 | 240 | 2 (0.8) | 2 | 0 | 0 | 0 | 2 (100) |

* ND, not determined.

**Figure S1. Five sequence clusters identified in this study.** Sequence types that have no more than 3 different loci with each other, but not assigned in MLST database were assigned to the same cluster. ST-5620 cluster included ST-5542, ST-5620, ST-10395, ST-11040, ST-11041, ST-11044 and ST-12877; ST-5662 cluster was composed of ST-3129, ST-5658, ST-5751, ST-8208, ST-8686, ST-9472, ST-9582, ST-9583, ST-9758, ST-9760, ST-10378, ST-10382, ST-10401, ST-10412, ST-10414, ST-10453, ST-11046, ST-12865, ST-12866 and 3 new sequence types with incomplete loci data (lack of *aroE* allele); ST-5819 cluster included ST-5819, ST-8788, ST-10108, ST-11042, ST-11043, ST-11047, ST-12861, ST-12864 and ST-12869; ST-5863 cluster was composed of ST-4830, ST-5636, ST-5863, ST-9587, ST-9588, ST-12860, ST-12862, ST-12867 and ST-12879; ST-9514 cluster included ST-4822, ST-9506, ST-9514, ST-9757 and ST-10398.

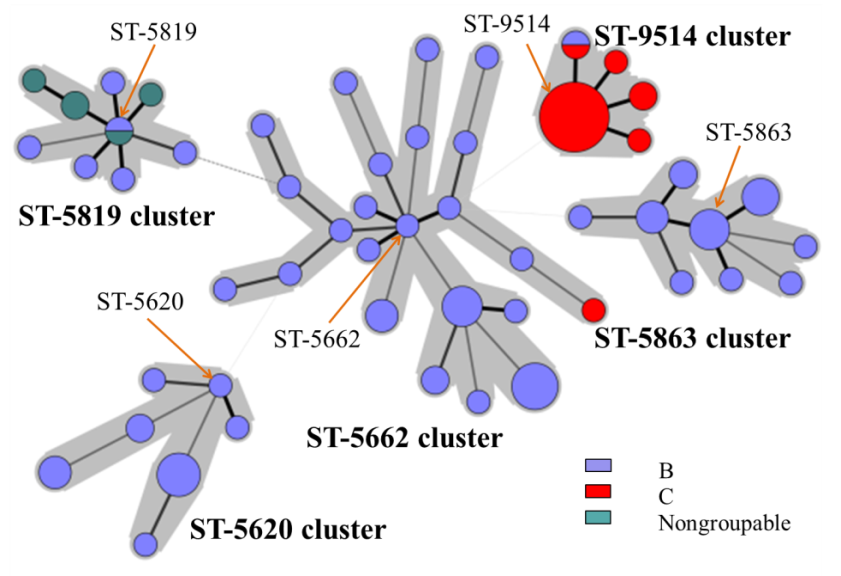
